# Conservation relevant fine scale distribution and habitat associations of threatened elasmobranchs in temperate nearshore waters

**DOI:** 10.1101/2025.06.30.662326

**Authors:** C.R. Hopkins, G. Cullen, R.L. Flatt, E.E. Brooker, D.M. Bailey, N.M. Burns

## Abstract

Elasmobranchs are globally threatened and experiencing ongoing declines. Understanding threatened elasmobranch distribution is critical for developing effective marine conservation strategies. However, our knowledge of fine scale elasmobranch habitat association and distribution in temperate nearshore systems is limited. Here, we examined the presence, relative abundance and habitat association of sharks, skates and rays using benthic baited remote underwater stereo-video systems (SBRUVs). From 772 deployments (682 total hours, 53 minutes average soak time) across three years (2021-2023) and two Scottish sea lochs and adjacent bays, elasmobranchs were detected on 31.2% of deployments (n = 241). Our surveys detected six species of elasmobranchs, representing 17.6% of the resident elasmobranch diversity reported to date in nearshore waters < 200 m depth in UK waters. The species detected include two species listed as globally Vulnerable on the IUCN Red List, spiny dogfish (*Squalus acanthias*) and porbeagle (*Lamna nasus*) and one Critically Endangered species, flapper skate (*Dipturus intermedius*). Critically Endangered flapper skate were detected in 5.2% deployments (n = 40) and were the only species recorded which did not show a relationship between the probability of presence and substratum type. Our findings provide critical data on the fine scale spatial distribution and habitat use of elasmobranchs, informing evidence-based conservation measures and supporting more consistent and targeted policy action for these species in Scotland.

## 1. Introduction

Globally, elasmobranchs are declining amid an ongoing extinction crisis (Dulvy et al. 2021, Pacoureau et al. 2021). Elasmobranch population declines have been documented at regional scales across the Mediterranean, the Caribbean, the eastern Red Sea and Australia (Ferretti et al. 2008, Ward-Paige et al. 2010, Spaet et al. 2016, Roff et al. 2018 respectively). Life history traits such as late maturity, low fecundity, and low intrinsic growth rates make elasmobranchs particularly sensitive to overexploitation (Stevens et al. 2000, Ward-Paige et al. 2012). Overexploitation is the key factor in the declines of oceanic sharks and rays (Dulvy et al. 2021), and is also driving high extinction risk in coral reef sharks and rays (Sherman et al. 2023).

In temperate coastal regions, elasmobranchs are often caught in multispecies fisheries or as bycatch, with many threatened elasmobranch species present in heavily fished temperate waters (Liu et al. 2022). In the Northeast Atlantic, these threatened species include the flapper skate (*Dipturus intermedius*), tope (*Galeorhinus galeus*), angelshark (*Squatina squatina*), spiny dogfish (*Squalus acanthias*), and porbeagle shark (*Lamna nasus*). The Northeast Atlantic and the Mediterranean account for nearly 12% (146 species) of described shark species, and waters around the UK show high elasmobranch functional rarity (rare traits combined with geographical restrictiveness) (Trindade-Santos et al. 2022, Coulon et al. 2023, Pimiento et al. 2023). Yet, the increasing extinction risk of elasmobranchs in the Northeast Atlantic and the Mediterranean is greater than any globally assessed vertebrate group on the Red List Index (RLI) (Walls and Dulvy 2021), making this region of high priority for elasmobranch conservation and management.

Elasmobranch conservation planning and fisheries management at the regional and national level is a priority (Stein et al. 2018). However, designing and implementing effective strategies is hindered by gaps in basic and applied research, including key data on species biology, abundance and population status, and scale and intensity of threats (Jorgensen et al. 2022). Batoids (skates, rays and guitarfish) in particular are under studied and accurate abundance estimates are needed (Sherman et al. 2018, de Oliveira et al. 2023). A better understanding of temperate elasmobranch habitat requirements and other factors that shape their distribution is also critical for their management and protection (Birkmanis et al. 2020, Espinoza et al. 2020a) as an improved knowledge of elasmobranch critical habitat across life stages can inform effective Marine Protected Area (MPA) design (Speed et al. 2010). Overcoming data gaps in elasmobranch research is logistically and financially challenging (Jorgensen et al. 2022), but advanced monitoring techniques including environmental DNA (eDNA), drone surveys and baited remote underwater video systems (BRUVS) are increasingly used to obtain fisheries independent data for this ecologically important group (Jorgensen et al. 2022, Liu et al. 2022).

The use of BRUVS to assess elasmobranch communities and abundance is growing (e.g. Osgood et al. 2019, Espinoza et al. 2020b, Bruns and Henderson 2020). BRUVS can provide a low impact fisheries independent method to sample populations of threatened elasmobranchs and are a good method in baseline and long-term studies, as images and videos can be georeferenced and kept as a permanent record to compare species compositions and abundances over temporal and spatial scales (Sih et al. 2017). Additionally, Stereo BRUV systems can enable precise measurements of body size which can be used to estimate maturity of individuals (Elliott et al. 2018). BRUVS can be deployed across a wide range of habitats and are not restricted to SCUBA depths and times (Whitmarsh et al. 2017).

However, the majority of BRUV surveys have focused on coral or rocky reefs, largely due to higher expected biodiversity on videos collected near reefs and improved visibility as a result of limited loose sediment (Whitmarsh et al. 2017). Use of BRUVS on unconsolidated high sediment temperate marine environments is limited but there are examples in temperate systems from fish assemblage and behaviour studies (e.g.Shea et al. 2020). Northeast Atlantic elasmobranch species that are nearshore and in depths less than 200m include many demersal species present across hard substrate and soft sediments (Ebert and Dando 2021). The use of BRUVS could allow us to detect a large proportion of elasmobranch diversity and collect habitat association data across varied substrata in coastal locations that are inaccessible using standard fisheries methods.

Here, we used SBRUVS to assess presence and relative abundance of elasmobranchs detected across a varied seascape. We aimed to (1) detect elasmobranch species and their relative abundance in two Scottish sea lochs and adjacent bays, Loch Eriboll and Little Loch Broom, on the north and west coasts of Scotland respectively, and (2) determine habitat associations of nearshore elasmobranchs in depths less than 200m using a Bayesian hierarchical spatial model framework.

## 2. Methods

### 2.1 Study area

From 2021 to 2023, we sampled two Scottish sea lochs and adjacent bays, Loch Eriboll (2021-2023) and Little Loch Broom (2022-2023) located on the north and west coasts of Scotland, UK respectively. Loch Eriboll is the only sea loch on Scotland’s north coast, experiences strong tidal currents and is exposed to wind and waves from the north and west. The average depth of the Loch Eriboll site is 27.76 m (± 20.33 m S.D.) and seabed sediments consist of gravels, sands, bedrock and soft sediment (Burns et al. 2024). Little Loch Broom is located within the West Highland Scottish marine region which consists of a mostly sheltered complex coastline with sea lochs, bays and islands and varying seabed sediments. The average depth of the Little Loch Broom site is 47.42 m (± 39.62 m S.D.) with boulder-strewn moraines and mud-rich basins across the seabed (Bradwell and Stoker 2023).

### 2.2 Data Collection

Benthic SBRUV surveys were carried out in Loch Eriboll in July and August 2021, 2022 and 2023 and in Little Loch Broom in 2022 and 2023. These SBRUV surveys were conducted as a component of a wider survey of benthic species and habitats across the north and west of Scotland. SBRUVS were deployed from small vessels during daylight hours (08:13 –17:52 hrs) at predefined random depth stratified stations in the two study locations. Three SeaGIS (https://www.seagis.com.au/) SBRUV frames were used. Two GoPro Hero 9 cameras per SBRUV frame were encased in individual SeaGIS waterproof housings set to video quality of 1080, super-wide frame and recording at 30 frames per second. Two dive torches per frame were attached to improve visibility. A bait canister containing approximately 500 g of chopped mackerel (*Scomber scombrus*) was placed 1.5 m at the cross-section in front of the cameras. SBRUVS were lowered from the boat on a rope with a surface marker buoy to facilitate retrieval, and each SBRUV deployment was left on the seabed for at least 50 minutes. SBRUVS were deployed at a minimum distance of 500 m from each other.

### 2.3 Video analysis

Video footage was reviewed by two observers; a subset of footage was independently reviewed by other co-authors to further verify species and substrate identification. The left camera’s footage always viewed when available, and if the left camera was obscured, footage from the right camera was viewed. The start time of the video was recorded when the SBRUV landed on the seafloor and ended when the SBRUV was recovered. For each video, deployment time, elasmobranch species presence, time of first sighting, and MaxN were recorded. MaxN, the maximum number of individuals of the same species appearing at the same time, was used to avoid repeat counts of individual animals re-entering the field of view (Cappo et al. 2003). The substratum type in each video was recorded using the categories developed in Burns et al. (2024). Elasmobranch species were identified to species level using morphological features. The total length of elasmobranch individuals appearing in the video frame with the highest MaxN for each species was measured using in EventMeasure software (SeaGIS-EM V5.51).

### 2.4 Statistical Analysis

All statistical analysis was conducted using R version 4.4.2 (R Core Team 2024), and with the R-INLA package for the modelling components (Lindgren and Rue 2015). We adopted a Bayesian hierarchical spatial model framework fitted in INLA for the presence of each of the elasmobranch species encountered at each of the two study sites (Loch Eriboll and Little Loch Broom). This hierarchical modelling approach has the advantage that the data observations and the underlying process which generated the data are to some extent modelled separately. The binary response variable was the presence or absence of individuals from the elasmobranch species modelled, mapped by a logit link function to the linear predictor. The linear predictor was composed of two parts, the fixed effect component and the spatial random field which can account for latent factors not explicitly included in the model, such as spatial dependencies. The Gaussian random field (GF) here is fitted using the Stochastic Partial Differential Equations (SPDE) approach to create the triangulation of the model domain (Fig 1.).

**Figure 1.**
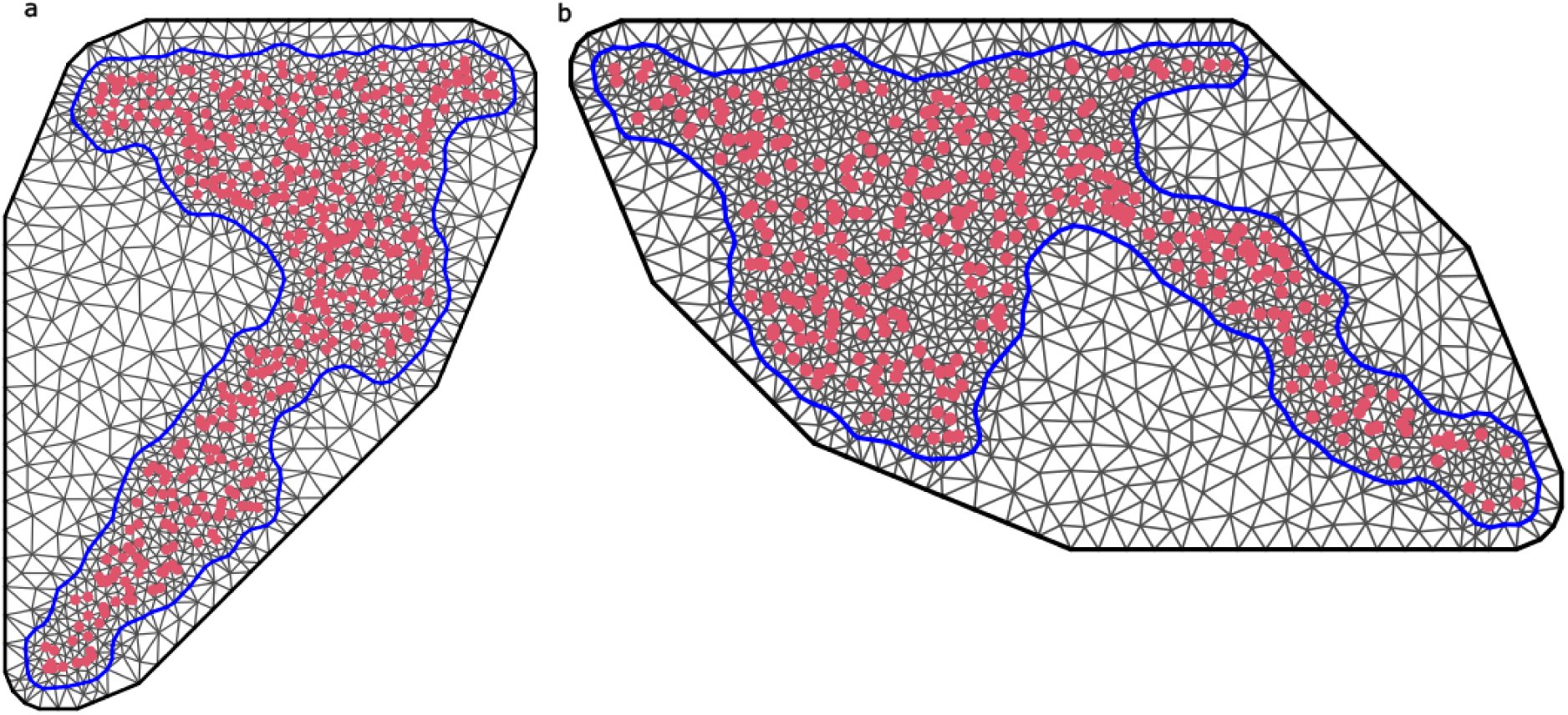
The mesh triangulation constructed for both the (a) Loch Eriboll and (b) Little Loch Broom study sites used in the SPDE approach. The blue line displays the mean low water line used as the boundary of the model domain and the red dots show the location of the SBRUV camera deployments.

We aimed to maintain the regular size and shape of triangle in the model domain for both sea lochs and extend the spatial extent of the triangulation to avoid boundary effects which can inflate the variance at the spatial boundaries (Lindgren et al. 2011). The triangulations obtained for LE contained 1,557 vertices using a maximum triangle edge length of 400 m in the model domain. The mesh obtained for the larger study area of LLB contained 1,989 vertices for the same 400 m maximum triangle edge length. The mean spring low water line was used to identify the model boundary domain. A backwards stepwise model selection process was carried out using WAIC and DIC to identify important variables and select between candidate models. During the selection process the spatial component in the models was always retained.

The default priors applied by R-INLA are flat normal, N (0, 31.62). Here, we fit a logistic model to the probability of elasmobranch presence, and if the default priors are used, once transformed through the logit-link the default flat normal prior piles the probability density at either end of the response scale (i.e. there are peaks at 0 and 1 with very low density in between). To avoid this we fitted weak sceptical priors to the models by prior predictive simulation (Appendix A. Fig 1.). We assumed that for species with an IUCN threatened conservation status, observations of these species were less likely and therefore were fitted with more sceptical priors.

## 3. Results

### 3.1 Elasmobranch species richness and diversity

A total of 785 benthic SBRUV deployments were made across the study period July-August 2021, 2022, and 2023 in the two study locations, Loch Eriboll and Little Loch Broom. Of the 785 deployments, 772 deployments were used in the analysis (Fig 2.); 13 were discounted due to the field of view being obscured. The total number of video hours was 688 hours with a minimum of 32 minutes, a maximum of 104 minutes, and an average time of 53 minutes (± 5.6 SD). All deployments were conducted during daylight hours from 08:13 hrs to 17:52 hrs. Deployment depths ranged from 0.5 m to 136.2 m, across both sites with a mean depth of 26.6 m (±18.3 m SD) in Loch Eriboll and 48.7 m (±31.6 m SD) in Little Loch Broom. Details of deployments based on location and survey period are provided in Table 2.

**Figure 2.**
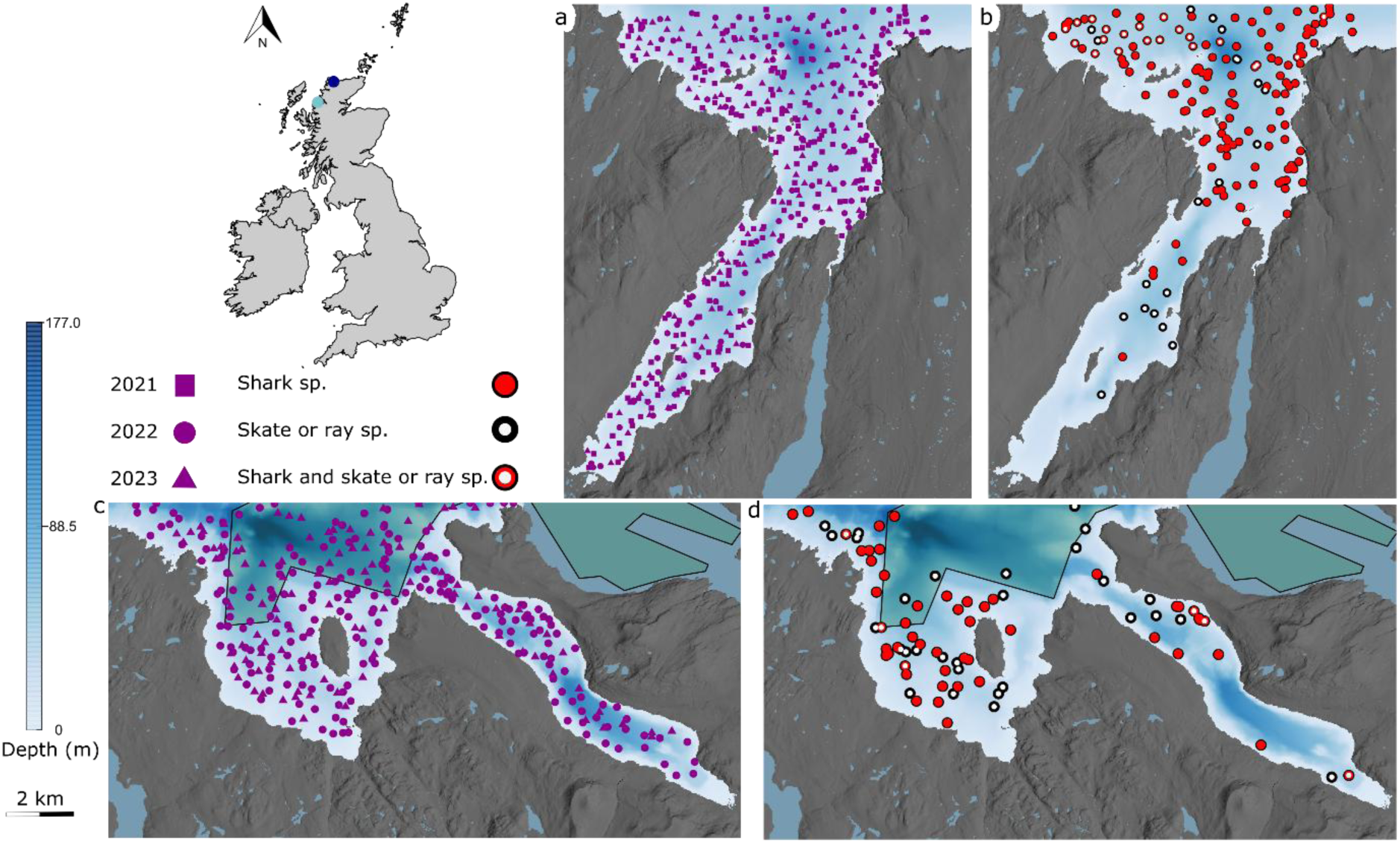
Map of the study areas in Scotland, UK with inset displaying the location of the study sites Loch Eriboll (dark blue circle) and Little Loch Broom (light blue circle). Pannels display (a) locations of SBRUV deployments from 2021, 2022 and 2023 for Loch Eriboll (n = 446), (b) locations of shark (red circles) and skate or ray (white circles) presence for Loch Eriboll (c) locations of SBRUV deployments in 2022 and 2023 for Little Loch Broom (n = 326) with permitted fishing trawl zone shown within the dark blue shaded area (n = 326) (d) locations of shark (red circles) and skate or ray (white circles) presence for Little Loch Broom.

**Table 2.**
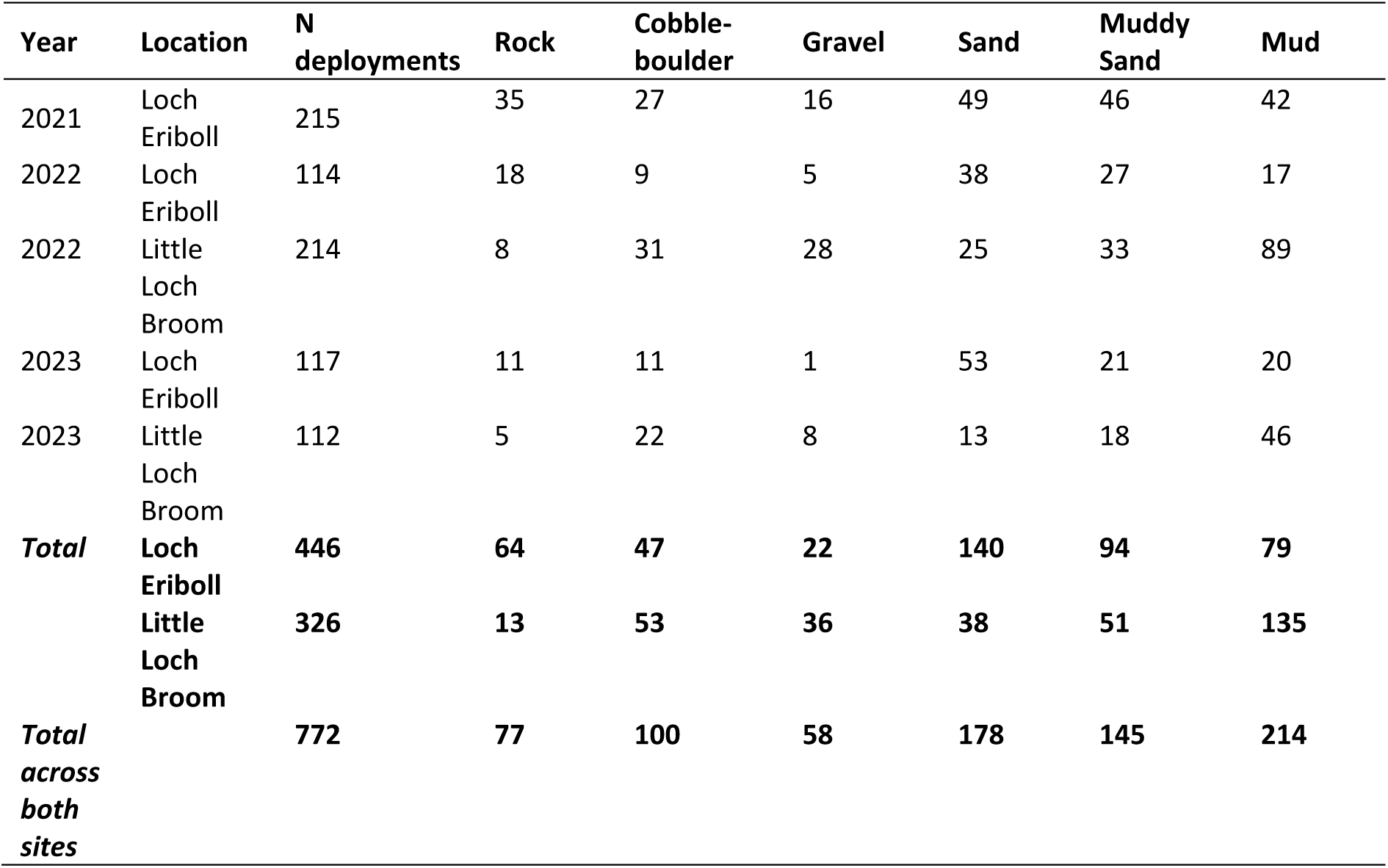
Summary of the number of Stereo Baited Remote Underwater Surveys (N deployments) completed in 2021, 2022 and 2023 by Location (Loch Eriboll and Little Loch Broom, Scotland), and Substratum type.

At both study locations and across all years and all deployments (n = 772), elasmobranchs were observed on 31.2% of the deployments (n = 241). The number of elasmobranch species recorded per SBRUV deployment varied from 1 to 3, and we detected a total of 6 elasmobranch species from 4 families and 5 genera (Fig 3. Table 3.). This is 17.6% of the resident elasmobranch diversity reported to date in nearshore Scottish waters < 200 m depth (14 resident shark species of which 3 species were observed and 20 skate and ray species of which 3 species were observed). A total of 6 elasmobranch species were detected in Loch Eriboll and 3 species detected in Little Loch Broom.

**Figure 3.**
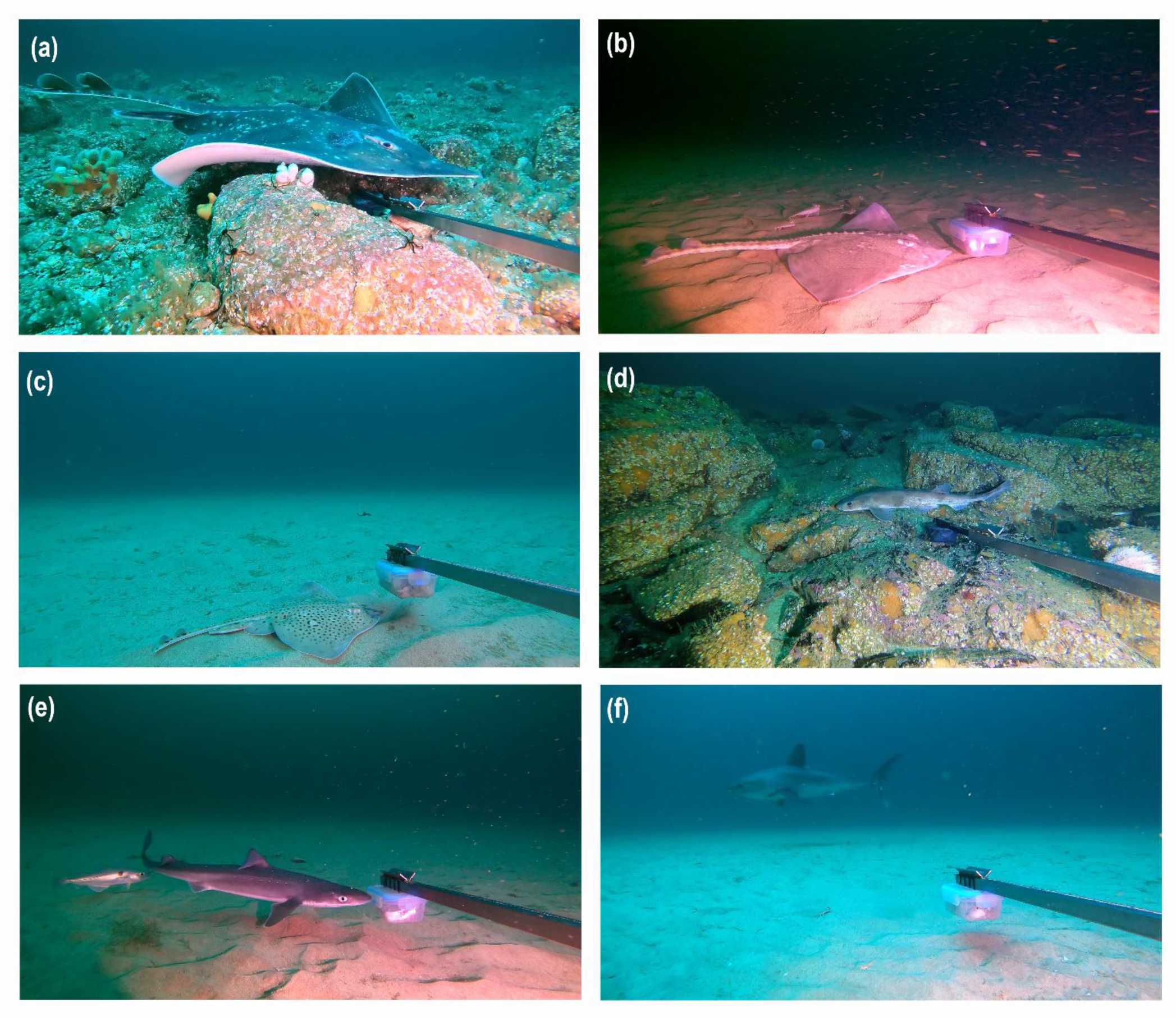
Images of elasmobranchs captured by the stereo baited remote underwater video systems (SBRUVS) in nearshore temperate waters < 200 m depth, Scotland, UK (a) flapper skate (*Dipturus intermedius*), (b) thornback ray (*Raja clavata*), (c) spotted ray (*Raja montagui*), (d) lesser spotted dogfish (*Scyliorhinus canicula*), (e) spiny dogfish (*Squalus acanthias*), (f) porbeagle (*Lamna nasus*).

**Table 3.** Summary of elasmobranch species recorded on Stereo Baited Remoted Underwater Video stations (SBRUVS) in Loch Eriboll (E) and Little Loch Broom (B), Scotland, % Occurrence from 772 total deployments (n = deployments with species present), size ranges (cm) as total length (TL), Conservation status (LC least concern, NT near threatened, VU vulnerable, and CR critically endangered) based on IUCN Red List status and population trend on the IUCN Red List (Version 2023-1).

| Common Name | Species Name | % Occurrence (n =<br>deployments with<br>species present) | MaxN | Size Range (cm) as<br>Total Length (TL) | IUCN Status | IUCN Population trend |
| --- | --- | --- | --- | --- | --- | --- |
| **SHARKS** |  |  |  |  |  |  |
| Lesser spotted<br>dogfish | *Scyliorhinus*<br>*canicula* | 24.4 (n = 188) | 3 | 34.3 – 88.0 | LC (Global); LC (Europe) | Stable (Global); Increasing (Europe) |
| Spiny dogfish<br>(spurdog) | *Squalus acanthias* | 0.9 (n = 7) | 2 | 56.5 - 87.3 | VU (Global); EN (Europe) | Decreasing (Global); Decreasing<br>(Europe) |
| Porbeagle | *Lamna nasus* | 0.1 (n = 1) | 1 | 239.4 | VU (Global); CR (Europe) | Decreasing (Global); Decreasing<br>(Europe) |
| **SKATES AND RAYS** |  |  |  |  |  |  |
| Flapper Skate | *Dipturus*<br>*intermedius* | 5.2 (n = 40) | 1 | 48.5 - 219.2 | CR (Global) | Decreasing (Global) |
| Thornback Ray | *Raja clavata* | 3.7 (n = 26) | 2 | 25.4 – 73.5 | NT (Global); NT (Europe) | Decreasing (Global); Stable<br>(Europe) |
| Spotted Ray | *Raja montagui* | 1.6 (n = 12) | 3 | 39.5- 63.4 | LC (Global), LC (Europe) | Increasing (Global); Stable (Europe) |

### 3.2 Habitat Association

Starting from a saturated model we performed backwards stepwise model selection using WAIC and DIC to identify influential variables (Table 4). The variable “Year” was included in the models for all species at both locations except for flapper skate which suggests fluctuation between years in the probability of observing lesser spotted dogfish, thornback ray and spotted ray, while flapper skate observations in these locations remained similar in each of the study years. The variable categorising the substratum type at the camera deployment sites was retained in all models except for flapper skate in Little Loch Broom. This indicates, for all the modelled species apart from flapper skate, the immediate seabed type, where the presence of the elasmobranch was recorded played a role in the probability of their presence.

**Table 4.**
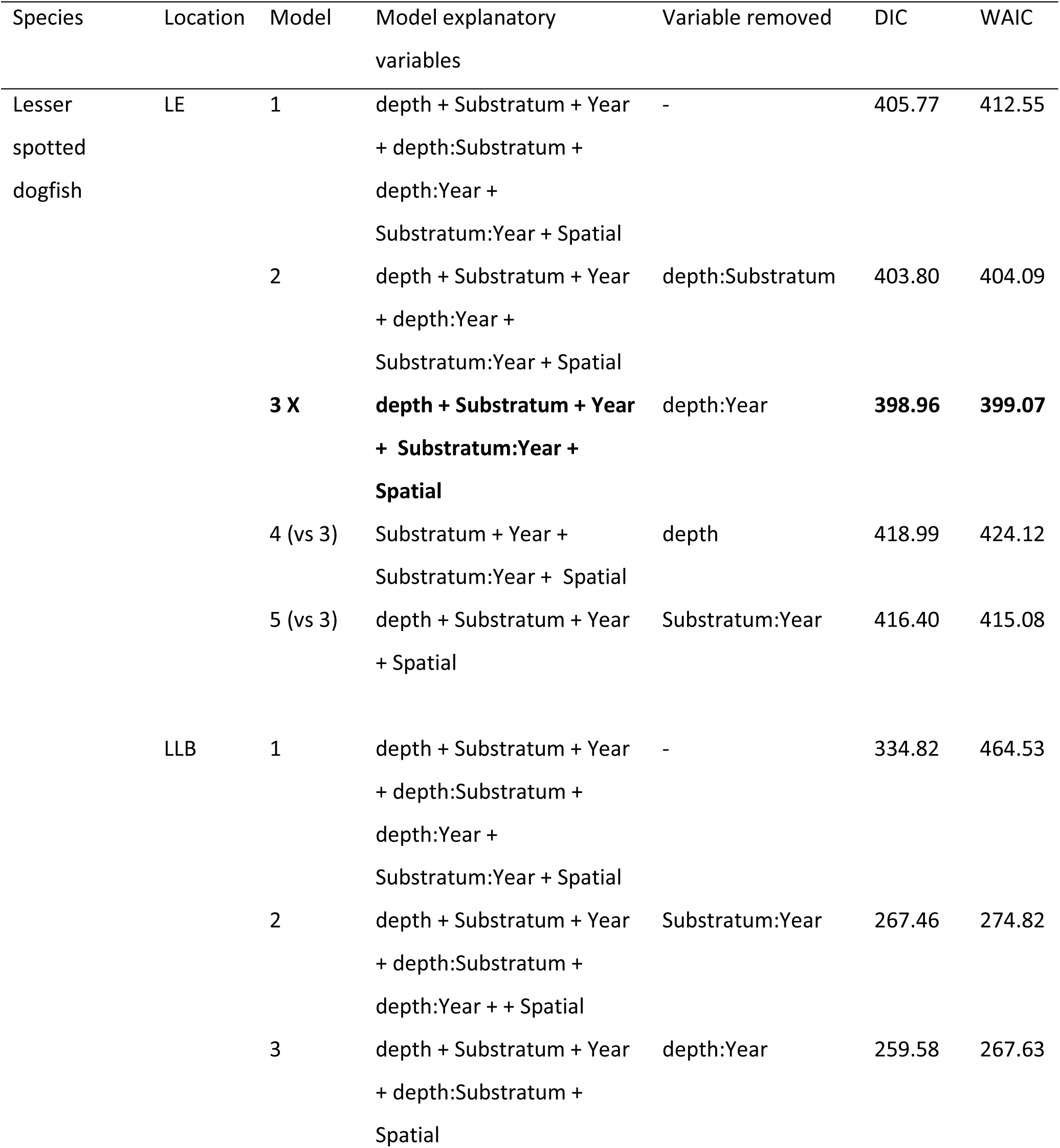
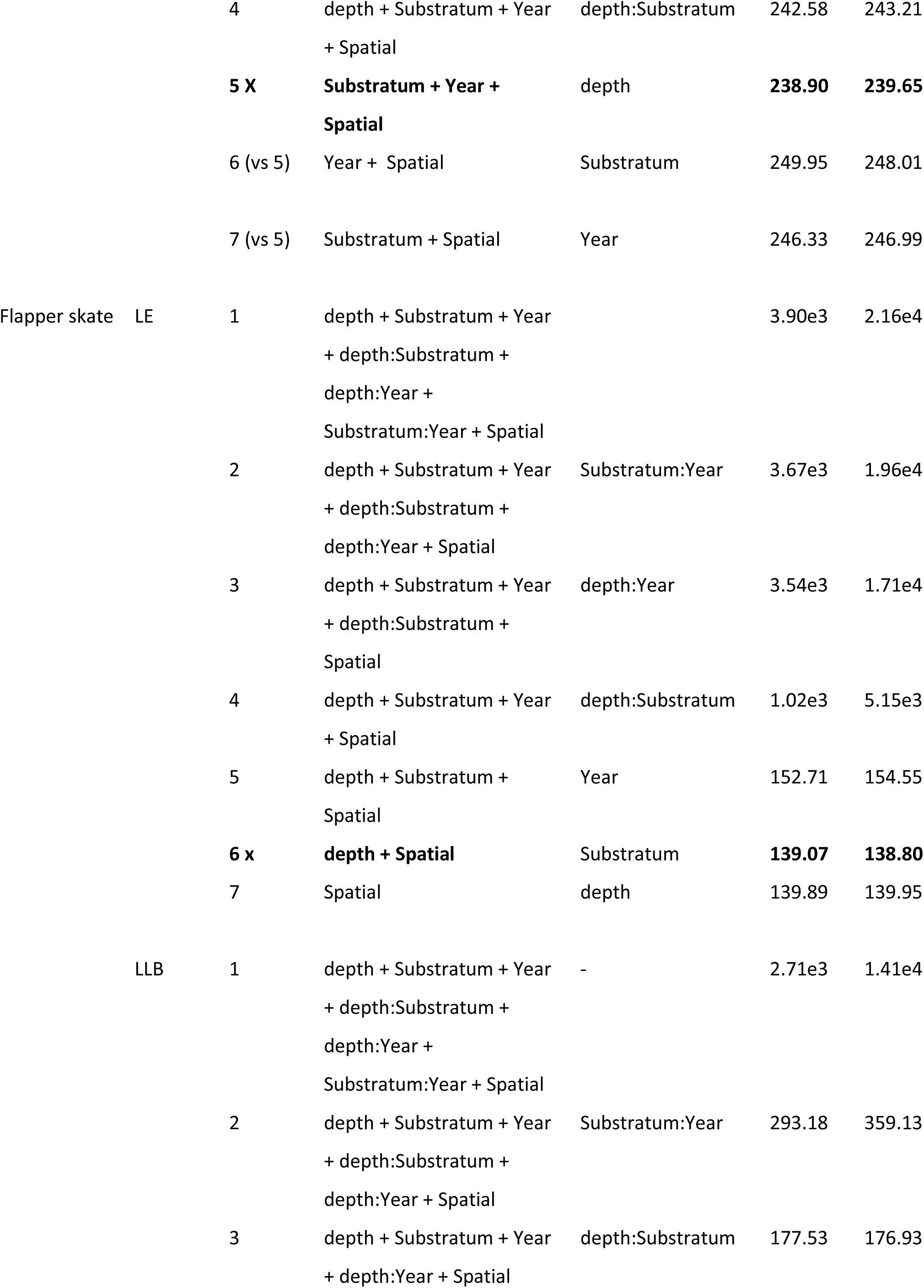
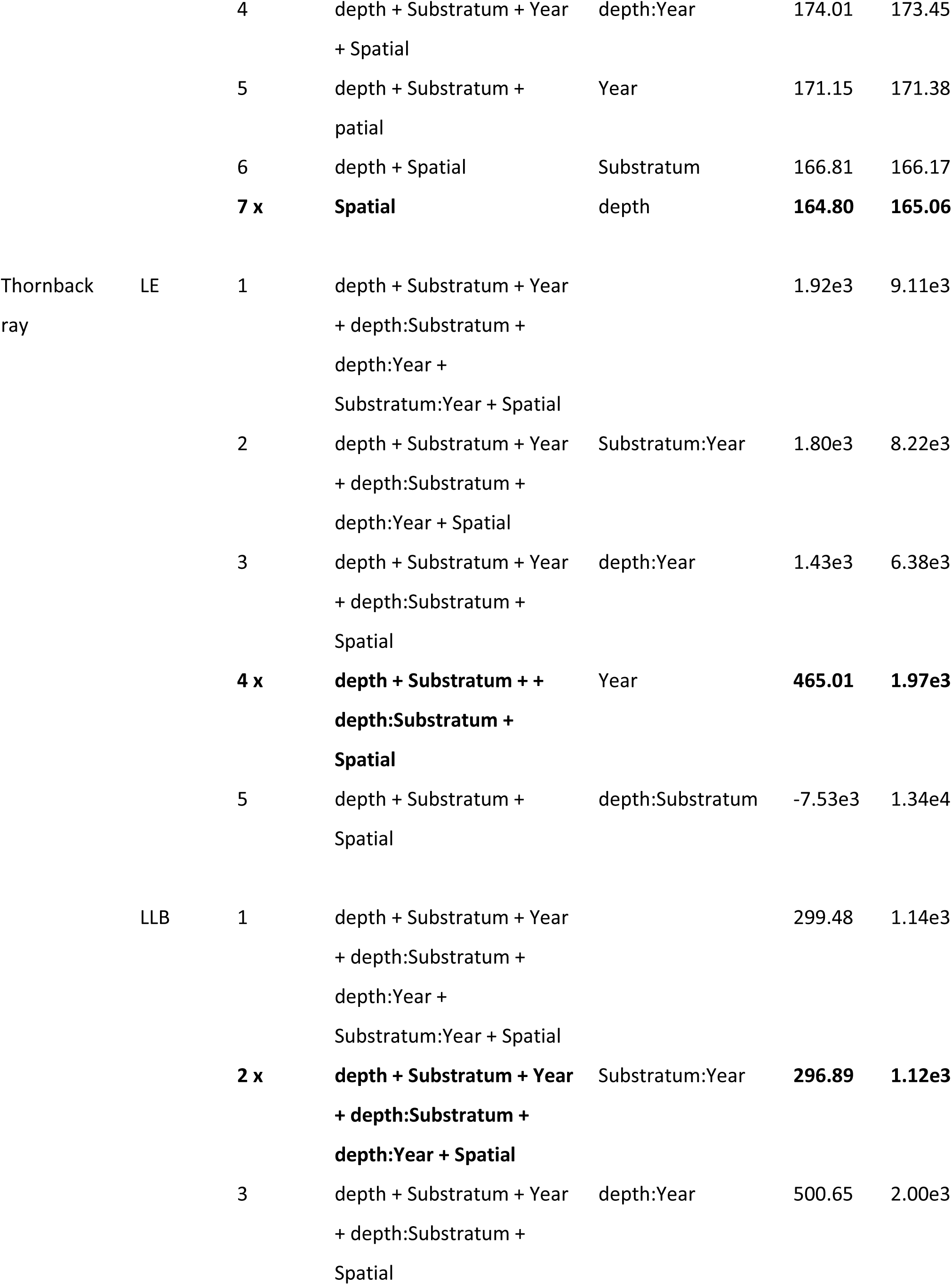
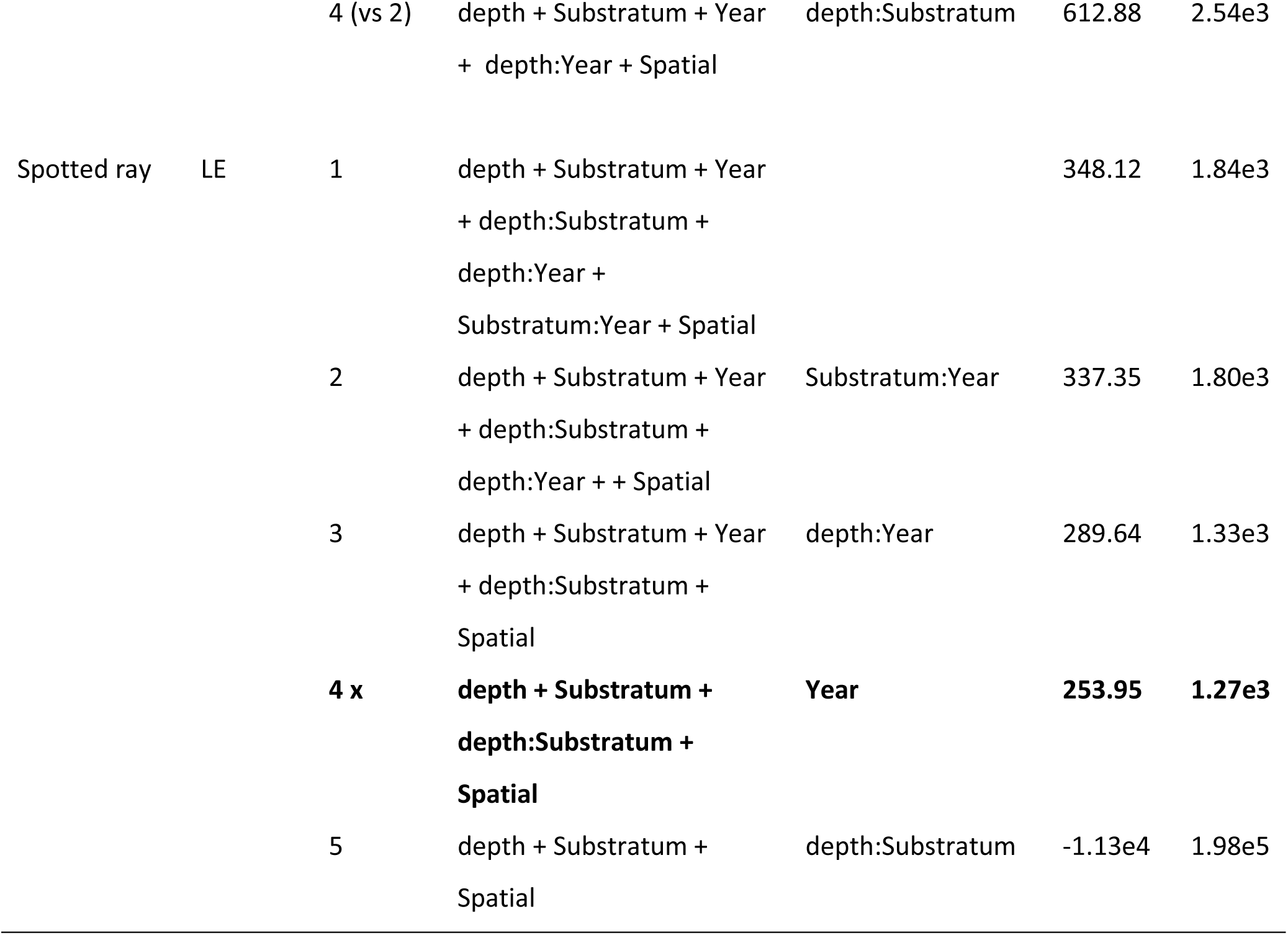
Model selection for INLA probability of species presence models for lesser spotted dogfish, flapper skate, and thornback ray in Loch Eriboll (LE) and Little loch Broom (LLB). Model selection for spotted ray only conducted for LE as none were observed in LLB. Backwards stepwise model selection was carried out using DIC and WAIC. Model comparisons should be assumed to be between sequential models in the list unless otherwise stated beside the model number. The optimal model is depicted by bold text and an **X** beside the model number.

| Species | Location | Model | Model explanatory variables | Variable removed | DIC | WAIC |
| --- | --- | --- | --- | --- | --- | --- |
| Lesser spotted dogfish | LE | 1 | depth + Substratum + Year<br>+ depth:Substratum +<br>depth:Year +<br>Substratum:Year + Spatial | - | 405.77 | 412.55 |
|  |  | 2 | depth + Substratum + Year<br>+ depth:Year +<br>Substratum:Year + Spatial | depth:Substratum | 403.80 | 404.09 |
|  |  | **3 X** | **depth + Substratum + Year**<br>**+ Substratum:Year +**<br>**Spatial** | depth:Year | **398.96** | **399.07** |
|  |  | 4 (vs 3) | Substratum + Year +<br>Substratum:Year + Spatial | depth | 418.99 | 424.12 |
|  |  | 5 (vs 3) | depth + Substratum + Year<br>+ Spatial | Substratum:Year | 416.40 | 415.08 |
|  | LLB | 1 | depth + Substratum + Year<br>+ depth:Substratum +<br>depth:Year +<br>Substratum:Year + Spatial | - | 334.82 | 464.53 |
|  |  | 2 | depth + Substratum + Year<br>+ depth:Substratum +<br>depth:Year + + Spatial | Substratum:Year | 267.46 | 274.82 |
|  |  | 3 | depth + Substratum + Year<br>+ depth:Substratum +<br>Spatial | depth:Year | 259.58 | 267.63 |
|  |  | 4 | depth + Substratum + Year<br>+ Spatial | depth:Substratum | 242.58 | 243.21 |
|  |  | **5 X** | **Substratum + Year +**<br>**Spatial** | depth | **238.90** | **239.65** |
|  |  | 6 (vs 5) | Year + Spatial | Substratum | 249.95 | 248.01 |
|  |  | 7 (vs 5) | Substratum + Spatial | Year | 246.33 | 246.99 |
| Flapper skate | LE | 1 | depth + Substratum + Year<br>+ depth:Substratum +<br>depth:Year +<br>Substratum:Year + Spatial |  | 3.90e3 | 2.16e4 |
|  |  | 2 | depth + Substratum + Year<br>+ depth:Substratum +<br>depth:Year + Spatial | Substratum:Year | 3.67e3 | 1.96e4 |
|  |  | 3 | depth + Substratum + Year<br>+ depth:Substratum +<br>Spatial | depth:Year | 3.54e3 | 1.71e4 |
|  |  | 4 | depth + Substratum + Year<br>+ Spatial | depth:Substratum | 1.02e3 | 5.15e3 |
|  |  | 5 | depth + Substratum +<br>Spatial | Year | 152.71 | 154.55 |
|  |  | **6 x** | **depth + Spatial** | Substratum | **139.07** | **138.80** |
|  |  | 7 | Spatial | depth | 139.89 | 139.95 |
|  | LLB | 1 | depth + Substratum + Year<br>+ depth:Substratum +<br>depth:Year +<br>Substratum:Year + Spatial | - | 2.71e3 | 1.41e4 |
|  |  | 2 | depth + Substratum + Year<br>+ depth:Substratum +<br>depth:Year + Spatial | Substratum:Year | 293.18 | 359.13 |
|  |  | 3 | depth + Substratum + Year<br>+ depth:Year + Spatial | depth:Substratum | 177.53 | 176.93 |
|  |  | 4 | depth + Substratum + Year<br>+ Spatial | depth:Year | 174.01 | 173.45 |
|  |  | 5 | depth + Substratum +<br>patial | Year | 171.15 | 171.38 |
|  |  | 6 | depth + Spatial | Substratum | 166.81 | 166.17 |
|  |  | **7 x** | **Spatial** | depth | **164.80** | **165.06** |
| Thornback<br>ray | LE | 1 | depth + Substratum + Year<br>+ depth:Substratum +<br>depth:Year +<br>Substratum:Year + Spatial |  | 1.92e3 | 9.11e3 |
|  |  | 2 | depth + Substratum + Year<br>+ depth:Substratum +<br>depth:Year + Spatial | Substratum:Year | 1.80e3 | 8.22e3 |
|  |  | 3 | depth + Substratum + Year<br>+ depth:Substratum +<br>Spatial | depth:Year | 1.43e3 | 6.38e3 |
|  |  | **4 x** | **depth + Substratum + +**<br>**depth:Substratum +**<br>**Spatial** | Year | **465.01** | **1.97e3** |
|  |  | 5 | depth + Substratum +<br>Spatial | depth:Substratum | -7.53e3 | 1.34e4 |
|  | LLB | 1 | depth + Substratum + Year<br>+ depth:Substratum +<br>depth:Year +<br>Substratum:Year + Spatial |  | 299.48 | 1.14e3 |
|  |  | **2 x** | **depth + Substratum + Year**<br>**+ depth:Substratum +**<br>**depth:Year + Spatial** | Substratum:Year | **296.89** | **1.12e3** |
|  |  | 3 | depth + Substratum + Year<br>+ depth:Substratum +<br>Spatial | depth:Year | 500.65 | 2.00e3 |
|  |  | 4 (vs 2) | depth + Substratum + Year<br>+ depth:Year + Spatial | depth:Substratum | 612.88 | 2.54e3 |
| Spotted ray | LE | 1 | depth + Substratum + Year<br>+ depth:Substratum +<br>depth:Year +<br>Substratum:Year + Spatial |  | 348.12 | 1.84e3 |
|  |  | 2 | depth + Substratum + Year<br>+ depth:Substratum +<br>depth:Year + + Spatial | Substratum:Year | 337.35 | 1.80e3 |
|  |  | 3 | depth + Substratum + Year<br>+ depth:Substratum +<br>Spatial | depth:Year | 289.64 | 1.33e3 |
|  |  | **4 x** | **depth + Substratum +**<br>**depth:Substratum +**<br>**Spatial** | **Year** | **253.95** | **1.27e3** |
|  |  | 5 | depth + Substratum +<br>Spatial | depth:Substratum | -1.13e4 | 1.98e5 |

Using the optimal model, we made predictions of the probability of presence for each species at both sites (Fig. 4). Lesser spotted dogfish had predicted probabilities of presence > 0 across all substrata in both Loch Eriboll and Little Loch Broom in at least one of the years. The lowest probabilities of presence were seen on softer substrata such as, mud and muddy sand. The probabilities predicted varied across all years of the study suggesting fluctuating abundancies. The prediction shown in Figure 4 b for Loch Eriboll included depth at a mean value of -26.8 m. As depth is increased the probability of presence tended to increase. Flapper skate were the only species recorded which did not show a relationship between the probability of presence and substratum type. In Little Loch Broom none of the fixed effects were retained, leaving only the spatial element of the model and only depth was retained in the model for Loch Eriboll. In Loch Eriboll, as depth increased the probability of presence of detecting flapper skate also increased (Figure 4 c). Thornback ray and spotted ray showed similar model trends (Figure 4 d, e & f). These species were only observed on a few substratum types; namely mud, muddy sand, and sand for thornback ray and spotted ray were only detected in Loch Eriboll on sand.

**Figure 4.**
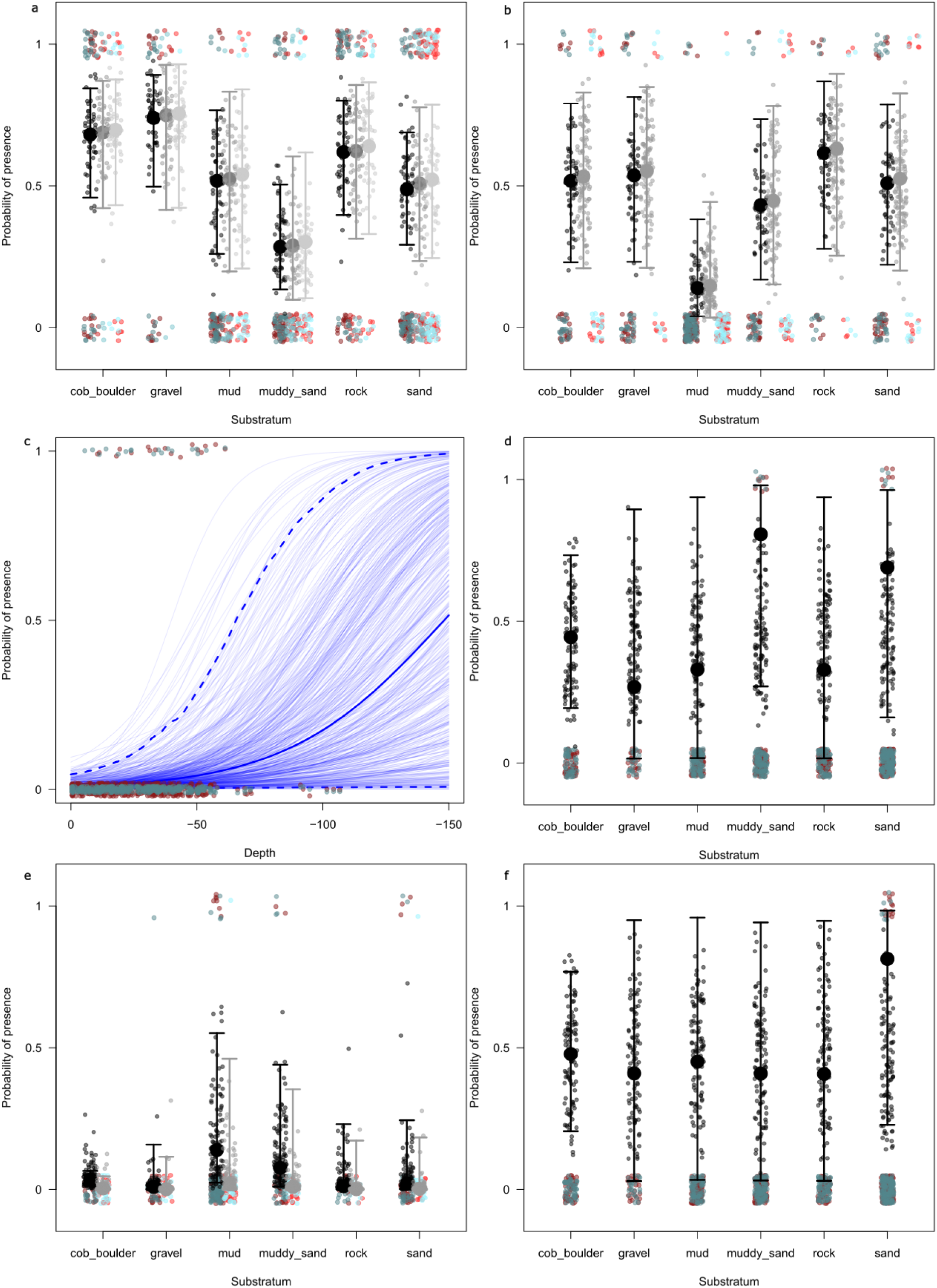
Model predictions displaying fixed effects for lesser spotted dogfish in Loch Eriboll (a) and Little Loch Broom (b), flapper skate in Loch Eriboll (c), thornback ray in Loch Eriboll (d) and Little Loch Broom (e), and spotted ray in Loch Eriboll (f). No fixed effects are reported here for flapper skate in Little Loch Broom as all the recorded variables dropped from the model during the selection process leaving only the spatial element of the model.

**Figure 5.**
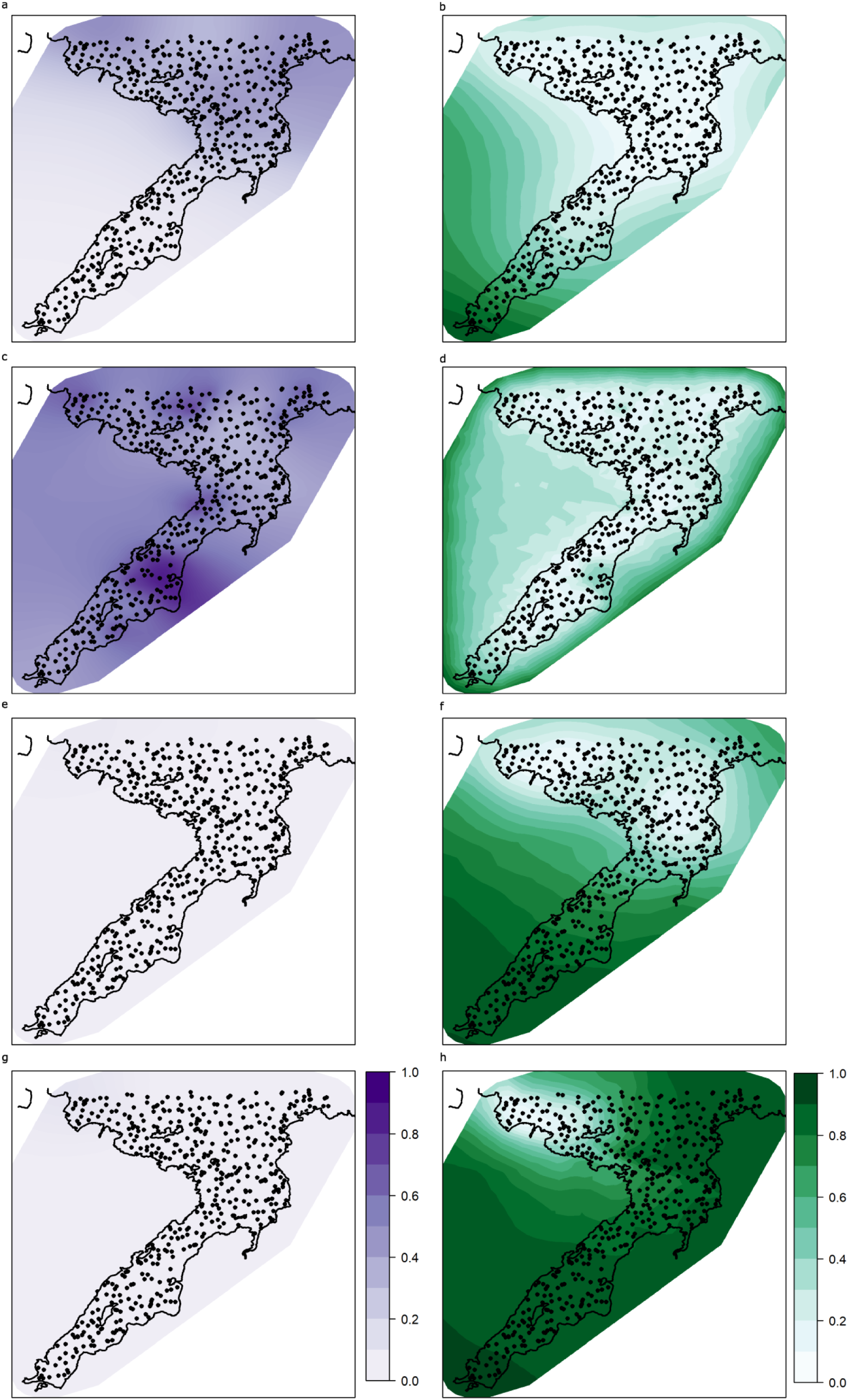
INLA Spatial random effect component of the presence absence models in Loch Eriboll for lesser spotted dog fish mean (a) and standard deviation (b), flapper skate mean (c) and standard deviation (d), thornback ray mean (e) and standard deviation (f), spotted ray mean (g) and standard deviation (h). The means and standard deviations are shown as inverse-link values, expressed on the response scale.

**Figure 6.**
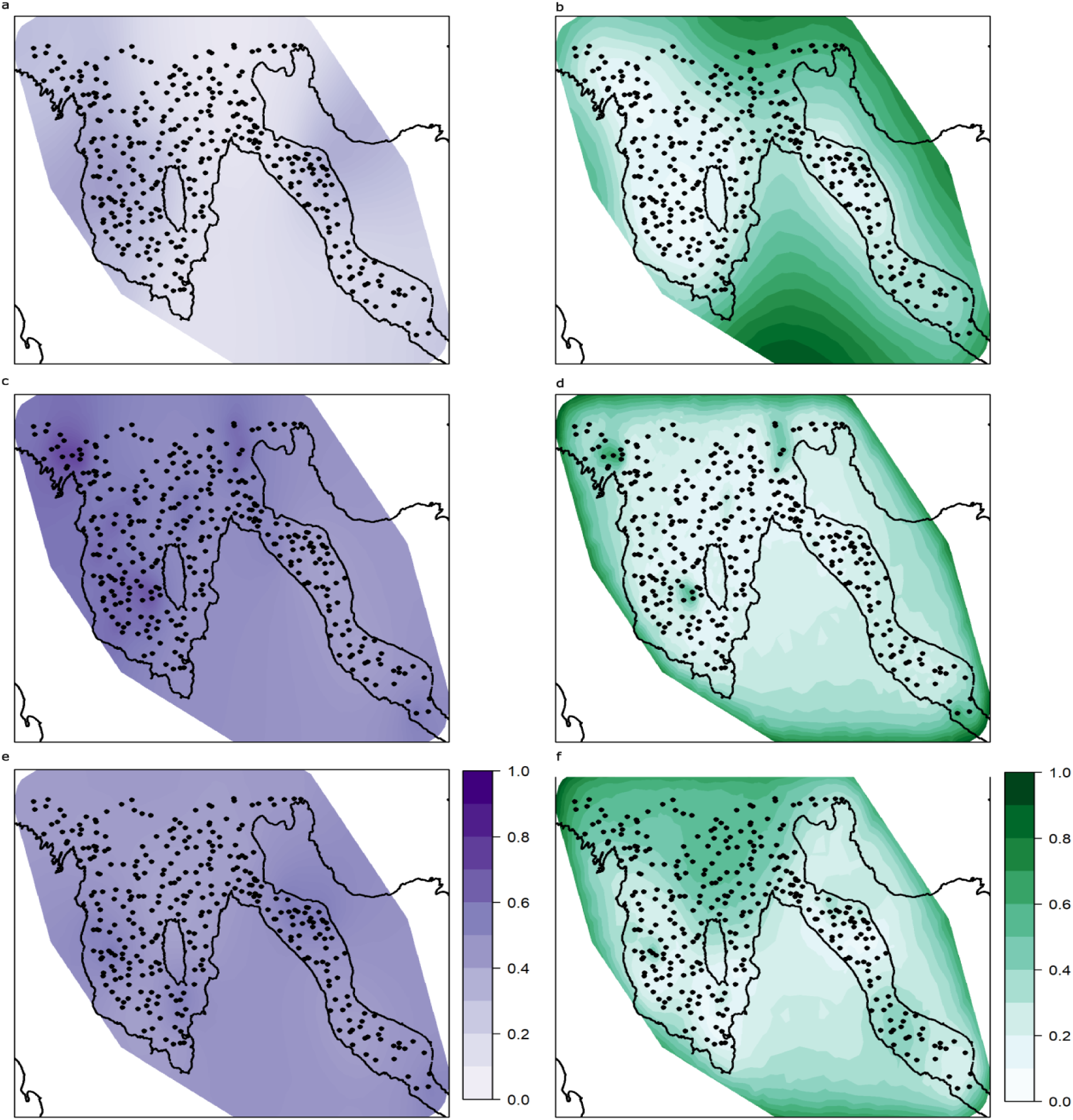
INLA Spatial random effect component of the presence absence models in Little Loch Broom for lesser spotted dog fish mean (a) and standard deviation (b), flapper skate mean (c) and standard deviation (d), thornback ray mean (e) and standard deviation (f). The means and standard deviations are shown as inverse-link values, expressed on the response scale.

## 4. Discussion

Using SBRUVS we have documented the presence of nearshore elasmobranch species across a range of depths and substratum types along the north and west coasts of Scotland. We detected six elasmobranch species in total, which is 17.6% of known resident nearshore Scottish elasmobranchs in depths less than 200 m (Ebert and Dando 2021). MaxN values were small for all species detected but elasmobranchs were detected on 31.2% of the deployments (n = 241). The most abundant elasmobranch species recorded in our study was the lesser spotted dogfish (*Scyliorhinus canicula*), a small, mesopredatory shark present in 24.4% of SBRUV deployments (n = 188). Larger elasmobranch species including the porbeagle shark (*Lamna nasus*) and flapper skate (*Dipturus intermedius*) were also detected. Flapper skate were the second most frequently observed elasmobranch at both survey locations, observed in 5.2% of deployments (n = 40). Previous studies on UK elasmobranchs indicate that threats such as overfishing and declines in habitat availability have contributed to low local abundance and species richness (Brander 1981, McHugh et al. 2011, Ellis 2016, Coulon et al. 2024), therefore our results are likely indicative of broader elasmobranch declines. However, here we used benthic SBRUVs that limit detection of pelagic species. We sampled during the day and in July and August (UK summer), therefore we were unable to observe diel changes in behaviour and activity of some species, and seasonal changes in community composition.

An understanding of habitat associations for individual elasmobranch species is essential for identifying important areas for elasmobranch conservation and to explain connectivity within and across ecosystems (Espinoza et al. 2014, 2020a, Jabado et al. 2018, Birkmanis et al. 2020). Due to the low detection of porbeagle and spiny dogfish (*Squalus acanthias*), having been observed in only one and seven SBRUV deployments respectively, it was difficult to identify spatial, or habitat association patterns for these species. Both species were only recorded on sandy habitat within the Loch Eriboll study site. Porbeagle were not expected to be observed in our study as this species exploits the water column, and here we used benthic SBRUVs. However, the porbeagle shark is also known to frequent northern Scottish waters, prey upon demersal fish and be sighted in the epipelagic zone (< 200 m) in summer months (Haugen and Papastamatiou 2019, Priede 2019, Ebert and Dando 2021). Captures of gravid porbeagle females with large embryos in Scottish waters suggest there may also be breeding grounds to the north (Gauld 1989, Priede 2019), but there is little information on how porbeagle may use north coast nearshore waters.

Lesser spotted dogfish are a generalist species associated with multiple habitats and here, our results suggest stronger associations across several seabed types predominantly comprised of harder substrata such as rock, cobble-boulder complexes and gravel. These seabed types are not suitable for trawling and therefore generally fall outside International Bottom Trawl Survey (IBTS) coverage, the data for which largely informs elasmobranch distribution models (Sguotti et al. 2016, McGeady et al. 2022). In Little Loch Broom, lesser spotted dogfish had the lowest probability of presence on mud, whereas in Loch Eriboll, the probability of presence was lowest on muddy sand. Thornback ray exhibited strong associations with sand and muddy sand in Loch Eriboll and mud, muddy sand and sand in Little Loch Broom. Spotted ray was observed and predicted to occur only on sand. There were no observations of either thornback ray or spotted ray on hard substrata and both species are known to forage, rest (bury) and deposit eggs in sandy habitats (Coulon et al., 2024; Nadia, 2024).

Flapper skate was the only species that did not demonstrate a strong association with any of the seabed substrata categories with the variable being dropped during model selection. Flapper skate are typically generalist in nature and are known to frequent multiple habitats (Dodd et al., 2022; Ellis et al., 2024). None of the fixed effect predictor variables were retained during model selection for the Little Loch Broom flapper skate model and, only depth was retained in the model for Loch Eriboll showing increased probability of presence in deeper water which is comparable to the likelihood of capture from angling studies (Benjamins et al. 2018). While little relationship was found between the habitat predictor variables and the probability of presence of flapper skate, this species displayed the highest mean parameter values in the spatial element of our models indicating clustering or local residency. Model selection also indicated no effect of year for either study site suggesting Loch Eriboll and Little Loch Broom could be sites of residency for flapper skate, which is an established movement pattern for the species (Lavender et al. 2022). While we did not observe strong associations between flapper skate and habitat in this study, strong habitat associations have been assumed to exist at key life stages, such as the association of flapper skate egg nurseries and rocky boulder habitat (Dodd et al. 2022).

While elasmobranchs are an increasing focus for fisheries management and marine conservation initiatives in the Northeast Atlantic, further co-operation and prioritisation for elasmobranchs at a European scale is recommended (Walls and Dulvy 2021). Of the six species observed in our study, commercial fishing is prohibited in EU and UK waters for two species (flapper skate and porbeagle) [Council Regulation (EU) 2020/123], two species (thornback ray and spotted ray) are currently regulated through a grouped common Total Allowable Catch (TAC), and lesser spotted dogfish have no TAC or minimum landing size in place (ICES 2022a, 2022b, 2023). The fishery for North Atlantic spiny dogfish reopened in 2023 with a total UK TAC of 7606 tonnes and only spiny dogfish under 100 cm are permitted to be landed [The Sea Fisheries (Amendment) Regulations 2023]. The UK Shark, Skate and Ray Conservation Plan (Defra 2011) developed to meet international obligations such as Convention on International Trade in Endangered Species (CITES) and Convention on the Conservation of Migratory Species of Wild Animals (CMS), aims to ensure the sustainable management of elasmobranch stocks by protecting critical habitat, introducing catch limits and increasing monitoring and reporting. There are considerable shortfalls to the implementation of the Plan, including lack of legislative force (the plan is non-statutory and largely advisory), lack of species-specific data, poor enforcement and limited integration into wider management frameworks (Defra 2013, McCully Phillips et al. 2015, Dulvy et al. 2017). There are proposals to strengthen measures for elasmobranch conservation in the Scottish Government’s Biodiversity Strategy Delivery Plan (Scottish Government 2023). However, availability of data on many elasmobranch species remains too poor to adapt or strengthen measures under changing pressures or environmental conditions (Delaval et al. 2022). Studies such as ours provide critical data on the fine scale distribution and habitat association of elasmobranchs, informing evidence-based conservation measures and supporting more consistent and targeted policy actions.

Here, we observed IUCN Red listed species that should be prioritised for targeted conservation measures and fisheries management. The porbeagle shark is on the OSPAR list of Threatened and Declining species, is globally Vulnerable, but Critically Endangered within Europe, and is in the order Lamniformes (mackerel sharks) which has a high evolutionary distinctness (Stein et al. 2018). Flapper skate and spiny dogfish are also both on the OSPAR list of Threatened and Declining species and are Critically Endangered and Endangered within Europe respectively. Porbeagle, spiny dogfish and flapper skate are Scottish Priority Marine Features (PMF), species of marine nature conservation importance in Scottish waters. However, only two Scottish Nature Conservation Marine Protected Areas (NCMPAs) currently have statutory measures in place to protect flapper skate, Loch Sunart to the Sound of Jura NCMPA (741 km^2^), and Red Rocks and Longay NCMPA (12.55 km^2^), with no NCMPAs in place for porbeagle or spiny dogfish.

The slow maturation and large size of flapper skate makes this species vulnerable to impacts from fishing gear and overexploitation throughout its life stages (Garbett et al. 2021, Régnier et al. 2021, Dodd et al. 2022). Within the Red Rocks and Longay NCMPA, prohibition of fishing and activities that disturb the seabed may help to preserve flapper skate egg nurseries (Dodd et al. 2022). However, demersal trawling outside these areas and MPA boundaries may contribute to bycatch of flapper skate (Lavender et al. 2022). Recent studies show limited recovery of flapper skate outside MPA boundaries highlighting that the present MPAs are not enough to facilitate wider population recovery (Régnier et al. 2024). Demersal trawling and dredging are prohibited in parts of the Little Loch Broom study site, situated within the Wester Ross NCMPA, but flapper skate is not a designated “feature” of the MPA. Flapper skate are a designated feature in only two NCMPAs across the whole of the Scottish MPA network; the lack of replication for flapper skate across the predicted range illustrates the lack of network connectivity planning for some species within the Scottish MPA network (Hopkins et al. 2016, 2020, Garbett et al. 2023). While there may be some incidental benefit for elasmobranch populations as a result of fisheries measures across the Scottish MPA network, any positive impact is unlikely to be detected by MPA monitoring where flapper skate and other elasmobranchs are not designated features and therefore are not subject to routine surveying. Here, we documented the presence of six species of elasmobranch within the Loch Eriboll study site, the site currently lacks elasmobranch specific conservation measures and there are no measures in place that protect benthic habitats or the water column which could offer incidental protection to elasmobranchs.

MPAs, when well managed and located to protect essential habitat, can contribute to elasmobranch conservation (Birkmanis et al. 2020). Areas with high species richness, areas where threatened species are present, or areas where there are unique species compositions are often targeted for MPA designation (Derrick et al. 2020). Here, we have used SBRUVS to identify areas where threatened elasmobranch species are present and associated with particular habitats not accessible to trawl surveys, which serves as valuable data for the design of protected elasmobranch sites.

However, while there is widespread advocacy for using MPAs to restore elasmobranch communities, reliably monitoring the abundance of elasmobranch species is needed to evaluate the effectiveness of such protection (Ward-Paige et al. 2012, Ward-Paige 2017, Liu et al. 2022). Understanding how elasmobranch species populations are connected across a seascape is also necessary to establish the most effective size and placement of replicate MPA sites to ensure protection across a species’ range (Conners et al. 2022, Gardner et al. 2024, Marraffini et al. 2024). Further research should focus on fine scale habitat association of elasmobranchs across the wider Scottish nearshore area and how this compares to MPA coverage in order to assess any gaps in elasmobranch protection in the midst of declining population trends.

## Acknowledgments

We thank Mr James Mather and Mr Louis Neate for assisting with field work, providing insight on the study locations, and skippering the research vessels. This study was funded in 2021 by Wildland Ltd. Through the Sustainable Inshore Fisheries Trust (SIFT). This work was further supported by the Scottish Government’s Rural & Environment Science & Analytical Services Division (RESAS), through their Strategic Research Programmes (2022–2027) to collect data from 2022. Bathymetric data is used courtesy of the Maritime and Coastguard Agency (MCA) and UK Hydrographic Office, collected as part of the INIS Hydro Interreg IVa Crossborder programme managed by the Special EU Programmes Body. Further funding support was provided by UKRI Natural Environment Research Council (NERC) Grant No. NE/X018342/1.

## Author contributions

CRH, NMB and DMB conceptualised this study. NMB and CRH acquired the funding for this project. NMB was in charge of the administration and field logistic coordination. NMB, CRH, GC and DMB collected the data in the field. CRH, GC, NMB, RF and EB completed the video analysis. NMB did all statistical analyses of the manuscript. CRH, NMB, RF and EB wrote the manuscript text. CRH prepared Figure 3 for publication. NMB prepared remaining figures for publication. All authors reviewed, edited and gave final approval for submission of the manuscript.

## Competing interests

The authors declare no competing interests.

## Data Availability Statement

All code and data used to produce the current manuscript are available at https://github.com/NeilMBurns/Temperate_Elasmobranchs and will have a DOI associated for publication through Fig Share after review.

## Ethical Statement

This study was observational in nature. No animals were captured, touched, or manipulated in any way. No animals were provisioned or consumed the bait. All observations were conducted in accordance with the ethical and animal welfare standards set by Scotland’s Rural College (SRUC).

## Supplementary material

**Table S1.** Prior specification used in final models. The priors specified here display the mean (*µ*) and standard deviation (*σ*). S.d. is presented here for simplicity, when specifying *σ* in INLA it is expressed as precision 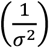. The flapper skate model for LLB only contained the spatial component and no spotted ray were observed in LLB.

| Species | LE models | LLB models |
| --- | --- | --- |
| Lesser spotted dogfish | Intercept N(0, 0.1) | Intercept N(0, 0.1) |
|  | Depth N(0, 0.01) | Substratum N(0, 1) |
|  | Substratum N(0, 1) | Year N(0, 0.1) |
|  | Year N(0, 0.1) |  |
|  | Year:Substratum N(0, 0.1) |  |
| Flapper skate | Intercept N(-5, 2) | - |
|  | Depth N(0, 0.15) |  |
| Thornback ray | Intercept N(-0.5, 0.05) | Intercept N(-0.5, 0.05) |
|  | Depth N(0, 0.05) | Depth N(0, 0.05) |
|  | Substratum N(0, 0.05) | Substratum N(0, 0.05) |
|  | Year N(0, 0.05) | Depth:Substaratum N(0, 0.05) |
|  | Depth:Substaratum N(0, 0.05) |  |
|  | Depth:Year N(0, 0.05) |  |
| Spotted ray | Intercept N(-0.5, 0.05) | - |
|  | Depth N(0, 0.05) |  |
|  | Substratum N(0, 0.05) |  |
|  | Depth:Substaratum N(0, 0.05) |  |

**Figure S1.**
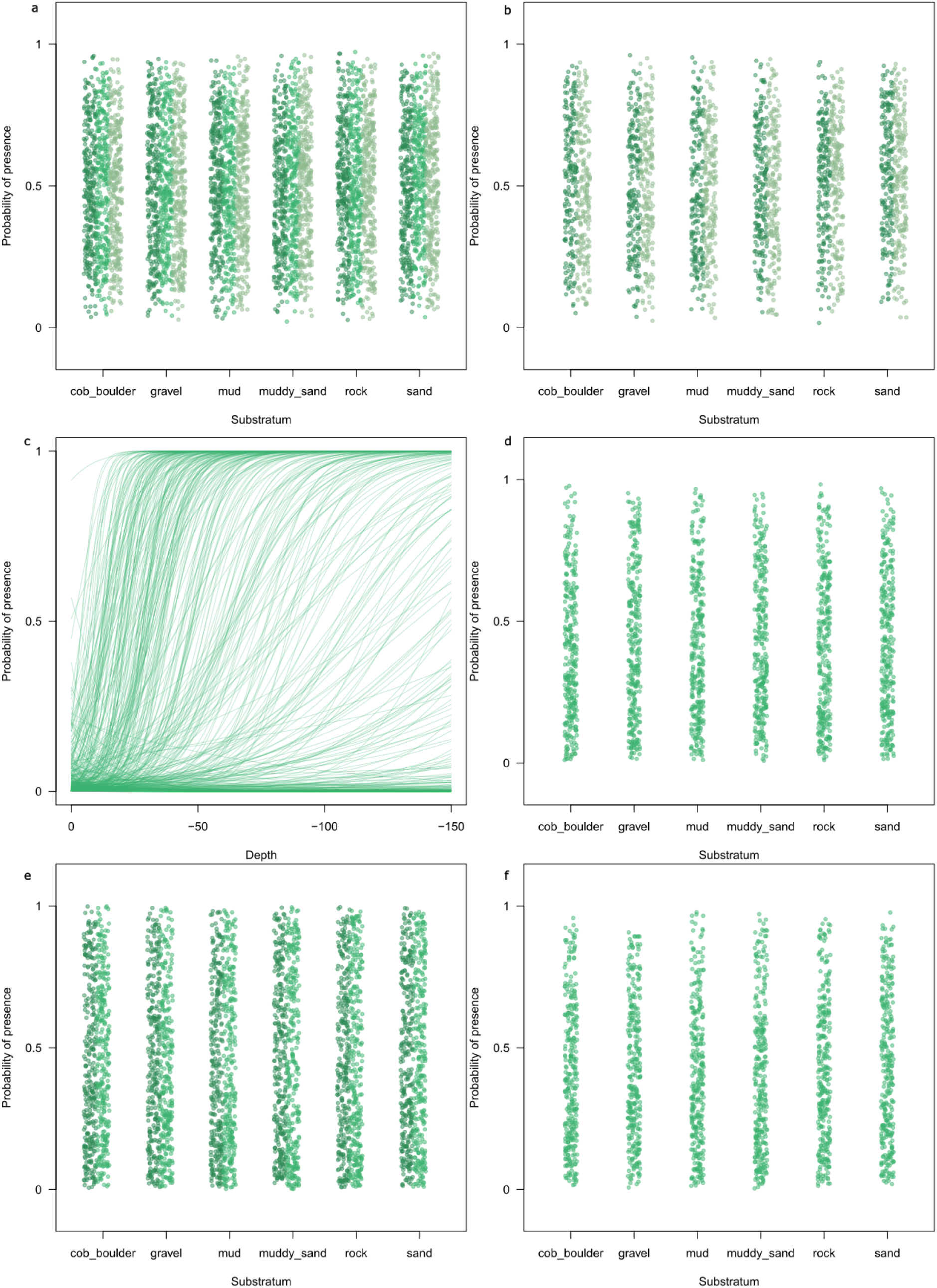
Prior predictive simulation for lesser spotted dogfish in Loch Eriboll (a) and Little Loch Broom (b), flapper skate in Loch Eriboll (c), thornback ray in Loch Eriboll (d) and Little Loch Broom (e), and spotted ray in Loch Eriboll (f). Green dots (and lines in (c)) indicate prior predicted means for 200 simulations in each category. Different shades of green dots correspond to different years.

## Notes

https://github.com/NeilMBurns/Temperate_Elasmobranchs

https://drive.google.com/drive/folders/1ncU-YkVfLp73J0rHNG9UgxnooocsIRr4?usp=drive_link

### Competing Interest Statement

The authors have declared no competing interest.

